# Single particle cryo-EM reconstruction of 52 kDa streptavidin at 3.2 Angstrom resolution

**DOI:** 10.1101/457861

**Authors:** Xiao Fan, Jia Wang, Xing Zhang, Zi Yang, Jin-Can Zhang, Lingyun Zhao, Hai-Lin Peng, Jianlin Lei, Hong-Wei Wang

**Affiliations:** Beijing Advanced Innovation Center for Structural Biology, School of Life Sciences, Tsinghua University, Beijing 100084, China; Tsinghua-Peking Joint Center for Life Sciences, Tsinghua University, Beijing 100084, China; Academy for Advanced Interdisciplinary Studies, Peking University, Beijing 100871, China

## Abstract

The fast development of single particle cryo-EM has made it more feasible to obtain the 3D structure of well-behaved macromolecules with molecular weight higher than 300 kDa at ~3 Å resolution. It remains a challenge to obtain high resolution structure of molecules smaller than 100 kDa using single particle cryo-EM, mainly due to the low contrast of the molecules embedded in vitreous ice. In this work, we applied the Cs-corrector-VPP coupled cryo-EM to study 52 kDa streptavidin (SA) protein supported on a thin layer of graphene film and embedded in vitreous ice. We were able to solve both the apo-SA and biotin-bound SA at near-atomic resolution using single particle cryo-EM. We demonstrated that the method is capable to determine the structure of molecule as small as 39 kDa and potentially even smaller molecules. Furthermore, we found that using the graphene film to avoid the adsorption to the air-water interface is critical to maintain the protein’s high-resolution structural information.

## Introduction

With the recent technical breakthroughs, cryo-EM has rapidly become one of the most powerful and efficient technology to investigate structures of macromolecules at near atomic resolution. In various cryo-EM structural determination methods, single particle analysis (SPA) draws most attention from structural biologists due to its relatively well-established methods in sample preparation, data collection, image processing and structural determination^1-3^. Thanks to the significant improvement on the recording speed and detective quantum efficiency (DQE) of the new direct electron detection cameras, more information at both low and high resolutions can be restored from raw movie stacks thus improving the reconstruction accuracy^4^. New algorithms based on Bayesian statistics have also greatly improved the efficiency of extracting signals from noisy micrographs and heterogeneous datasets^5-9^. Nowadays, it has become more and more routine to get the reconstruction of a well behaved protein with molecular weight larger than 300 kDa at ~3 Å resolution. In contrast, it remains a challenge to solve high resolution structure of proteins with molecular weight smaller than 200 kDa using SPA Cryo-EM. The major hurdle lies in the weak contrast of small-sized molecules embedded in vitreous ice under the conventional transmission electron microscopy (CTEM). Another major obstacle remaining in SPA cryo-EM is the adsorption of proteins to the air-water interface of the thin layer of solution during the cryo-specimen preparation^3,10,11^. This is more severe for small proteins because the suspended vitreous ice needs to be thinner than that embedding larger macromolecules in order to enhance the signal-to-noise ratio of the cryo-EM images. Until now, the smallest protein solved by CTEM using SPA at near atomic resolution is the 3.8?Å resolution structure of the 93?kDa isocitrate dehydrogenase (IDH)^12^.

In recent hardware developments, new apparatuses are introduced to cryo-EM, including energy filter, Cs-corrector and volta phase plate (VPP), to further improve imaging quality. VPP can introduce an extra phase shift to the contrast transfer function (CTF) of the objective lens thus increasing the low frequency signal of weak phase objects such as the frozen-hydrated biological molecules^13-15^. With new algorithms supporting the CTF determination and correction of micrographs taken with VPP^6,16,17^, it has been proved that VPP can be used to study various structures at near atomic resolution, including the 64 kDa hemoglobin at 3.2 Å resolution^18-21^. Using a combination of VPP and Cs-corrector, we demonstrated that the structure of apo-ferritin can be solved at near atomic resolution in both under- and over-focus modes of the objective lens^22^.

In this work, we used SPA cryo-EM with VPP and Cs-corrector to determine the structure of SA with a molecular weight of about 52 kDa. Different from hemoglobin which consists mostly of α-helices, SA constitutes of mainly β-strands, therefore is more difficult for precise alignment of the single particle images. Our work demonstrated that VPP renders SPA with the capability to solve SA in both apo-state and biotin-bound state at near atomic resolution. The biotin ligand can be distinguished clearly in the EM density. We also found that the graphene film can serve as a good supporting film to keep the small-sized SA in its intactness for high resolution structural determination. Our results proved in principle the capability of SPA cryo-EM in solving atomic models of small-sized proteins and their ligand-bound complexes. This would be of potentially great relevance in structural-based drug discovery.

## Results

### Preparation of frozen-hydrated SA specimen on graphene supporting film

In this study, we used the single-crystalline monolayer graphene over Quantifoil R0.6/1 gold grid as supporting film to facilitate cryogenic SA specimen preparation (See Materials and Methods for more details). Using a modified version of our previous imaging strategy to combine Cs-corrector and VPP for cryo-EM, we were able to collect high-resolution dataset of vitrified SA specimens with high efficiency (Materials and Methods). Examined under the VPP-Cs-corrector-coupled Titan Krios at 300 kV with phase shift ranging from 30 to 120 degrees, the SA specimens demonstrated monodispersed particles with high contrast that could be easily identified and automatically picked (Figure 1A). We found that the single-crystalline graphene with monolayer of carbon atoms introduced very low background noise to the specimen and could also serve as a good reference for the assessment of the cryo-EM image quality and motion correction with its hexagonal lattice signal^23,24^. After motion correction of the raw movie stacks of the specimen, we calculated the Fourier transform of the motion-corrected micrographs. In micrographs with good quality, we could observe clear reflection spots at 2.13 Å resolution in a hexagonal pattern corresponding to the graphene’s lattice (Figure S1), indicating a successful motion correction with high-resolution information recovered to at least 2.13 Å. It is worth noting that these reflection spots were not clear or sharp enough without the proper motion correction (Figure S1). Therefore, the sharpness of the reflection spots of single-crystalline graphene in the Fourier transform could serve as a good indicator to judge the quality of the micrographs and the motion-correction efficiency.

**Figure 1.**
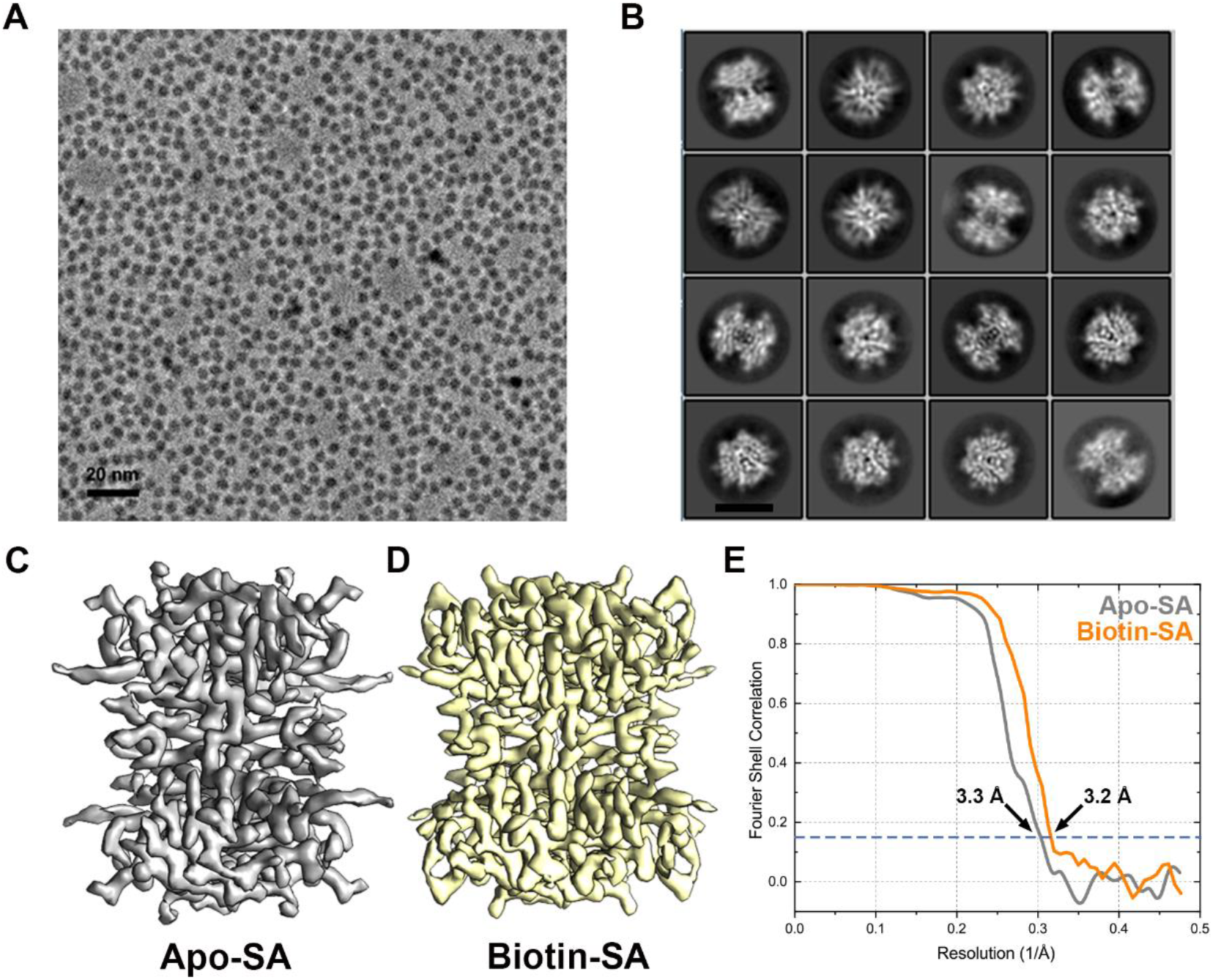
The SPA cryo-EM of SA. (A) A representative micrograph of SA specimen by VPP-Cs-corrector coupled cryo-EM. (B) Representative 2D class averages of SA particle images. The scale bar represents 5 nm. (C) The 3D reconstruction of apo-SA at 3.3 Angstrom resolution from 23,991 particles, and (D) the 3D reconstruction of biotin-bound SA at 3.2 Angstrom resolution from 45,686 particles. (E) The FSC curves of the two reconstructions using gold standard criteria.

### Single particle reconstruction of SA by VPP-cryo-EM

Using the automatic particle picking algorithm in Relion software^5^, we extracted about 710,000 and 1,350,000 particle images from the good motion-corrected micrographs of SA in the absence and presence of biotin, respectively, and applied a 120 Å Fourier high-pass filter to the particles prior further processing (Figure S2). The high-pass filter turned out to be necessary for correct alignment of the particle images (Figure S2), in agreement to our previous results^25^. Reference-free two-dimensional (2D) alignment and classification from such dataset yielded 2D class averages with clear secondary structural features that matched with the atomic model of SA protein (Figure S2B and S2C). Using an initial model generated *de novo* using Stochastic Gradient Descent (SGD) method in Relion, we performed multiple rounds of three dimensional (3D) classifications to screen the best particles for final 3D refinement and reconstruction (Figure S3). We obtained a reconstruction of apo-SA at 3.3 Å resolution (with D2 symmetry applied during the refinement, Figure 1C) from a final dataset composed of ~25,000 particles and a reconstruction of SA-biotin complex at 3.2 Å resolution (with D2 symmetry applied during the refinement, Figure 1D) from a final dataset comprising of ~45,000 particles, respectively, as validated by the gold standard Fourier shell correlation (FSC, Figure 1E). We also performed reconstructions of the two different states without imposing any symmetry (Figure S4A). The reconstructions have very similar map quality to those calculated with D2 symmetry, albeit slightly lower resolutions (Figure S4C).

The 3D reconstructions of SA in its apo- and biotin-bound state were both clear enough to depict all the secondary structural elements and visualize most of the side chains (Figure 2 and 3, Movie S1). The atomic model of SA solved previously by X-ray crystallography (PDB 1MEP ^26^) can fit into the EM densities with correlation coefficient ~0.74, indicating the structural fidelity of SA in its crystallographic and soluble forms. The density of biotin in the SA-biotin reconstruction can be precisely identified with the unambiguous docking of biotin’s atomic model (Figure 2). Compared with the biotin-bound SA, the density corresponding to loop 46-51 in the EM map of apo-SA was missing (Figure 2), indicating that this “lid-like” loop is flexible without ligand binding. In contrast, this loop can be clearly defined in the EM map of biotin-bound SA and the major side chains (ASN23, SER27, TYR43, ASN49, SER88) that form stable hydrogen bond network around the biotin ligand are better resolved (Figure 2).

**Figure 2.**
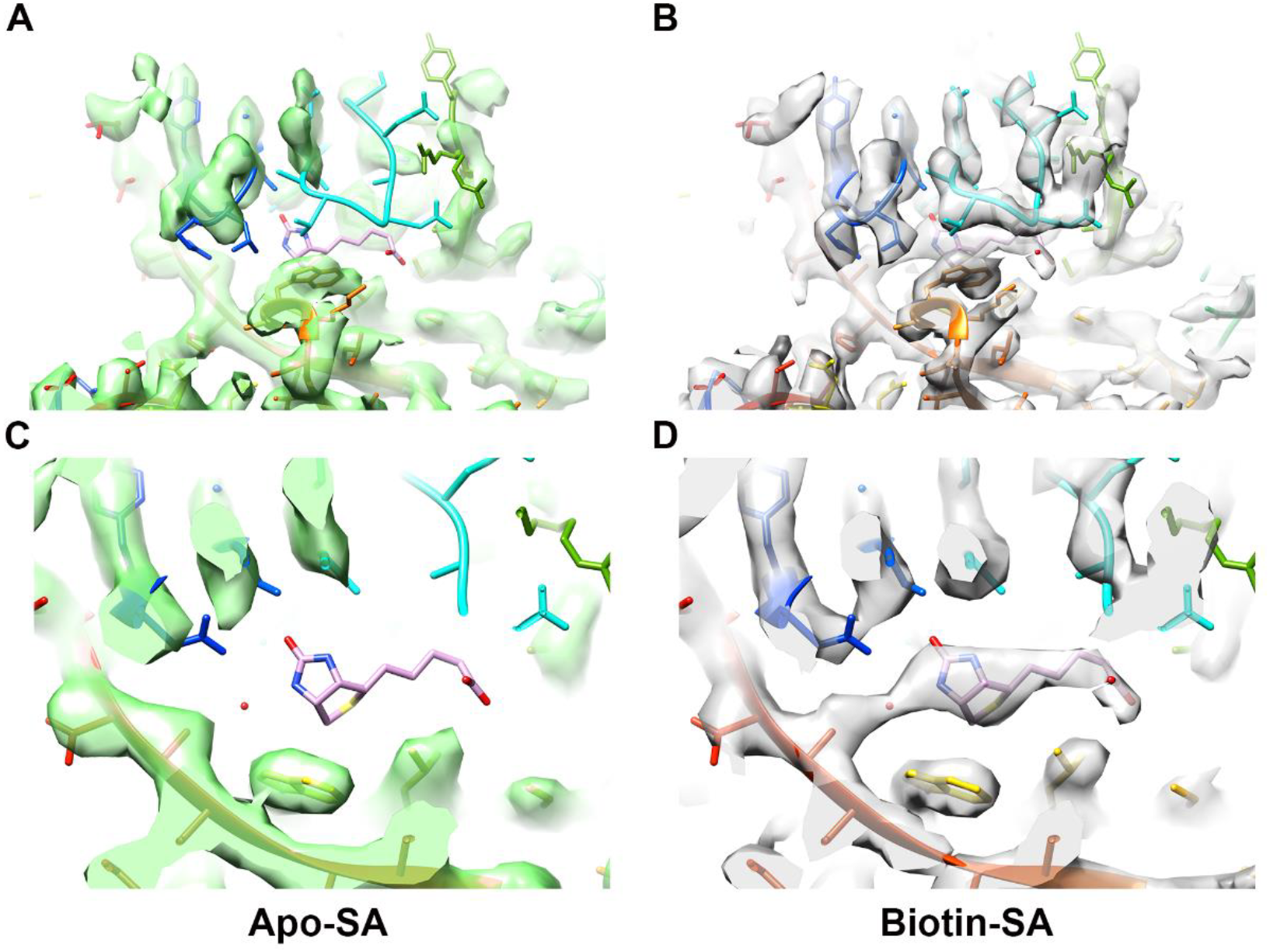
Comparison between the reconstructions of two SA states. (A and C) The region around the biotin-binding pocket of the apo-SA EM map has an empty density of the pocket and a missing loop 46-51 density, while in (B and D) the biotin-bound SA EM map these two densities are well resolved with the atomic model of biotin ligand and the loop 46-51.

**Figure 3.**
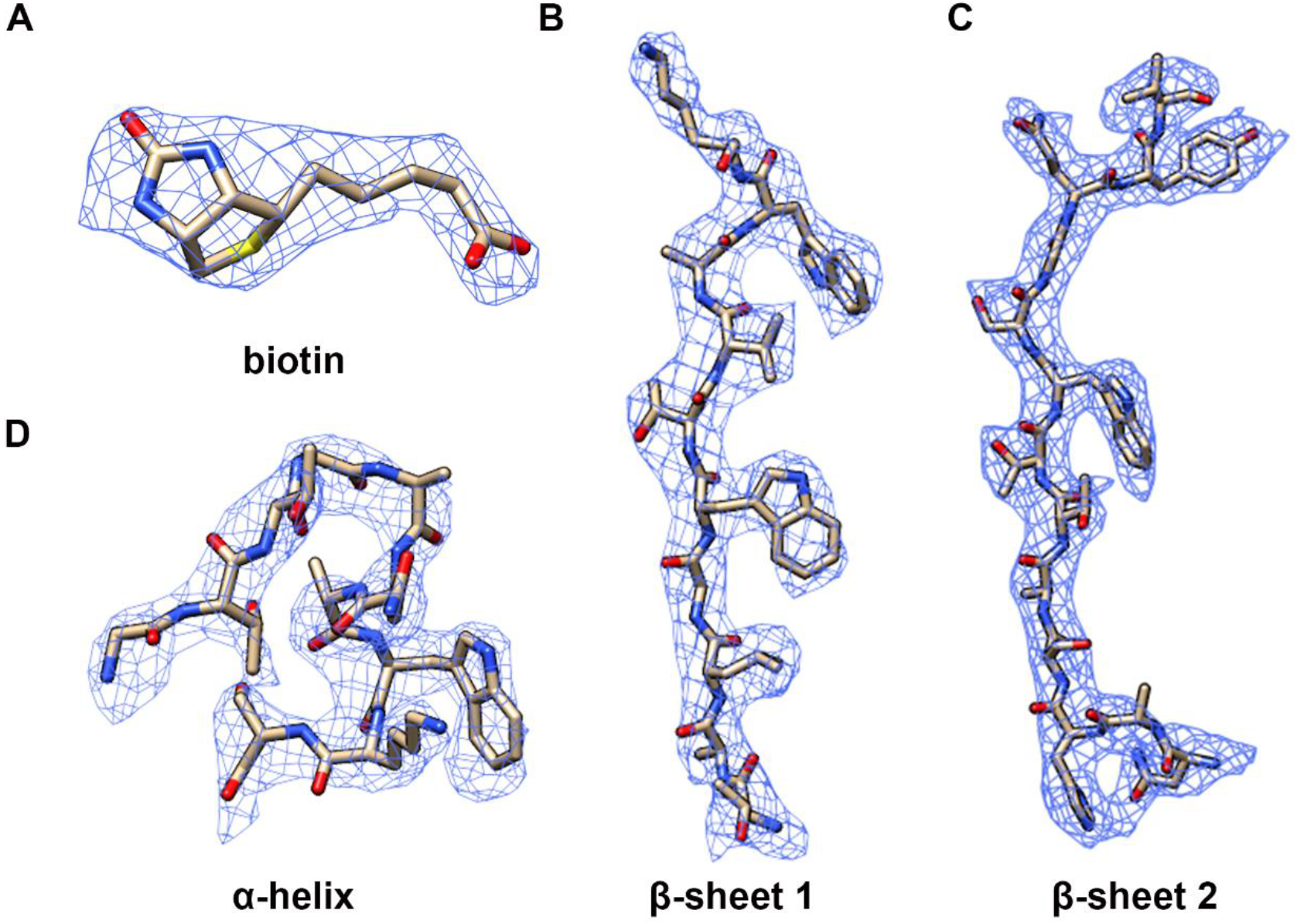
Representative regions of the biotin-SA EM map with their corresponding atomic models. (A) Biotin density in the binding pocket. (B) Representative densities of secondary structures: β-sheet (B and C) and α-helix (D).

### Focused classification analysis of the biotin binding pocket of SA

A critical problem of drug discovery is to identify the ligand binding site of the target protein. We wondered if ligand binding site can be determined via image processing in small proteins such as SA without a prior knowledge. Because SA is a tetramer and has four biotin binding sites in each protein, we treated each SA monomer (with one binding pocket) as an asymmetric unit and used the angular information from the reconstruction with D2 symmetry to align the four asymmetric units from the same particle to a given orientation. This step generated a three times larger dataset comprising roughly aligned asymmetric particles, thus called asymmetric particle dataset. After a local search refinement with C1 symmetry, the asymmetric particle dataset was subjected to 35 iterations of 3D classification into 4 classes in a skip-alignment mode in Relion. Without specific focusing on the binding pocket, a soft mask slightly larger than SA monomer was applied in either refinement or classification. We performed this 3D skip-alignment classification analysis of the apo-SA and biotin-SA datasets separately and found rather small occupancy variance around the biotin-binding pocket among different classes in each dataset (Figure S5), demonstrating unambiguously the lack of biotin in all the monomers of apo-SA and fully occupancy of biotin in all the monomers of biotin-SA. This is because SA has a very strong binding affinity to biotin and the condition of the biotin-SA specimen allowed full occupancy of the protein’s ligand binding sites. For other proteins and other conditions, the ligand occupancy may not be full. Thus, we tried to test whether we could extract the binding information by image processing from particles with partial ligand binding. We mixed the apo-SA and biotin-bound SA datasets and analyzed them as one dataset for 3D refinement. The reconstruction of the mixed dataset demonstrated a structure with biotin-like density in the binding pocket at 3.1 Å resolution (Figure 4A, 4B, S4B). From this mixed dataset of apo-SA and biotin-SA, a 3D skip-alignment classification of asymmetric unit into four classes illustrated distinct difference in the biotin binding pocket with or without biotin density (Figure 4C). While Class II was vacant of biotin density, all the other three classes all had biotin in the binding pocket. The results above proved the capability of heterogeneity analysis for ligand binding detection of small proteins by single particle cryo-EM.

**Figure 4.**
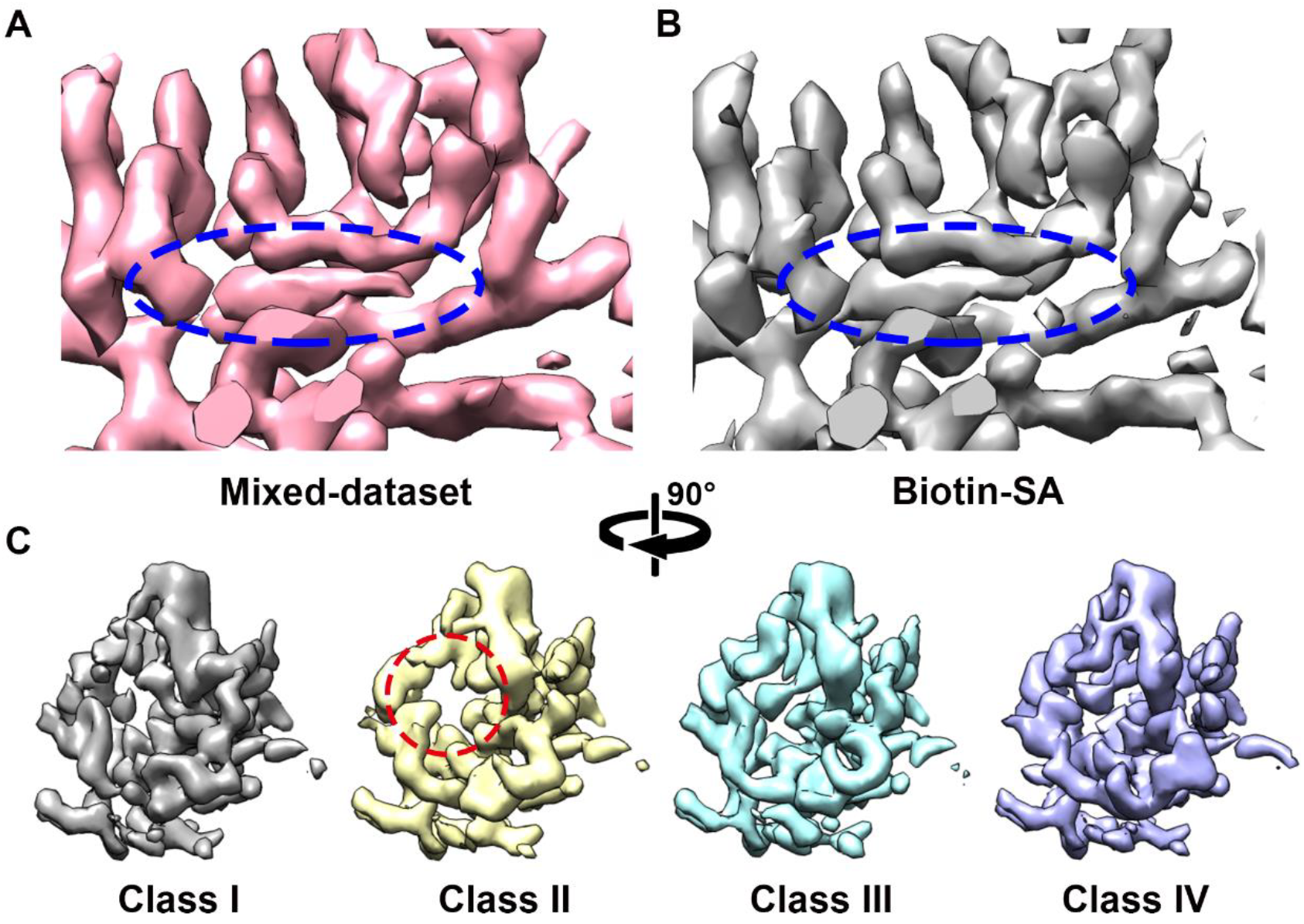
Reconstruction and classification using the asymmetric mixed-dataset. (A) The 3D reconstruction of the asymmetric mixed-dataset (apo-SA + biotin-SA) with 3.1 Angstrom resolution demonstrates a biotin-bound-like density (circled in blue) as (B) the 3D reconstruction of biotin-SA at 3.2 Angstrom resolution of a monomer. (C) Asymmetric 3D classification reconstructions of the mixed-dataset. The empty density of biotin-binding pocket in the monomer of Class II reconstruction is circled in red in contrast to the ligand densities in other classes.

### Reconstruction of sub-tetrameric SA from subtracted dataset

While the 52 kDa SA is the smallest protein solved at near atomic resolution using SPA cryo-EM until now, we were wondering if SPA cryo-EM is capable to reconstruct even smaller proteins. We used the particle segmentation and subtraction algorithms^27,28^ that is currently available in Relion to generate monomeric (13 kDa), dimeric (26 kDa) and trimeric (39 kDa) SA datasets from raw biotin-SA datasets *in silica* (Figure 5A). The subtracted SA datasets had smaller molecular weights and broke the intrinsic D2 symmetry of SA, therefore the signal for proper alignment is even weaker. To verify whether the subtracted datasets can still generate valid 3D reconstructions, we used the roughly correct angular information to perform local 3D refinement. The angular information of each subtracted particle was calculated in accordance to the relative orientation in the original tetrameric SA particle and the angular information of that tetrameric SA image in the final tetramer reconstructions. Indeed, the reconstructions using given angular information were correct for all three subtracted datasets (Figure 5C). We further tested if the images from those three datasets had enough signal for searching the correct angular information without given any pre-knowledge. The 39 kDa trimeric dataset had enough signals to generate a correct 3D refinement result by global angular search from scratch (Figure 5C). By contrast, the monomeric and dimeric datasets failed to reconstruct from scratch (Figure 5C), verifying the lack of sufficient signals for global angular search in them. Besides, 2D classification of the three datasets using the given correct angular information without alignment all generated good 2D class averages with the correct shapes and features (Figure 5B, Skip Align). By removing all the angular information, we performed reference-free 2D alignment and classification of the three datasets from scratch in Relion. In this procedure, the 13 kDa monomeric dataset generated 2D class averages with roughly correct outlines but much noisier features than the perfectly aligned controls in different views (Figure 5B, Search Align, left panel), suggesting more alignment error in the reference-free alignment. The 26 kDa dimeric dataset generated one well-aligned view (Figure 5B, Search Align, middle panel), while the other views were misaligned alike the monomeric dataset. The 39 kDa trimeric dataset generated correct shapes and features in different views (Figure 5B, right panel), indicating a precise reference-free alignment.

**Figure 5.**
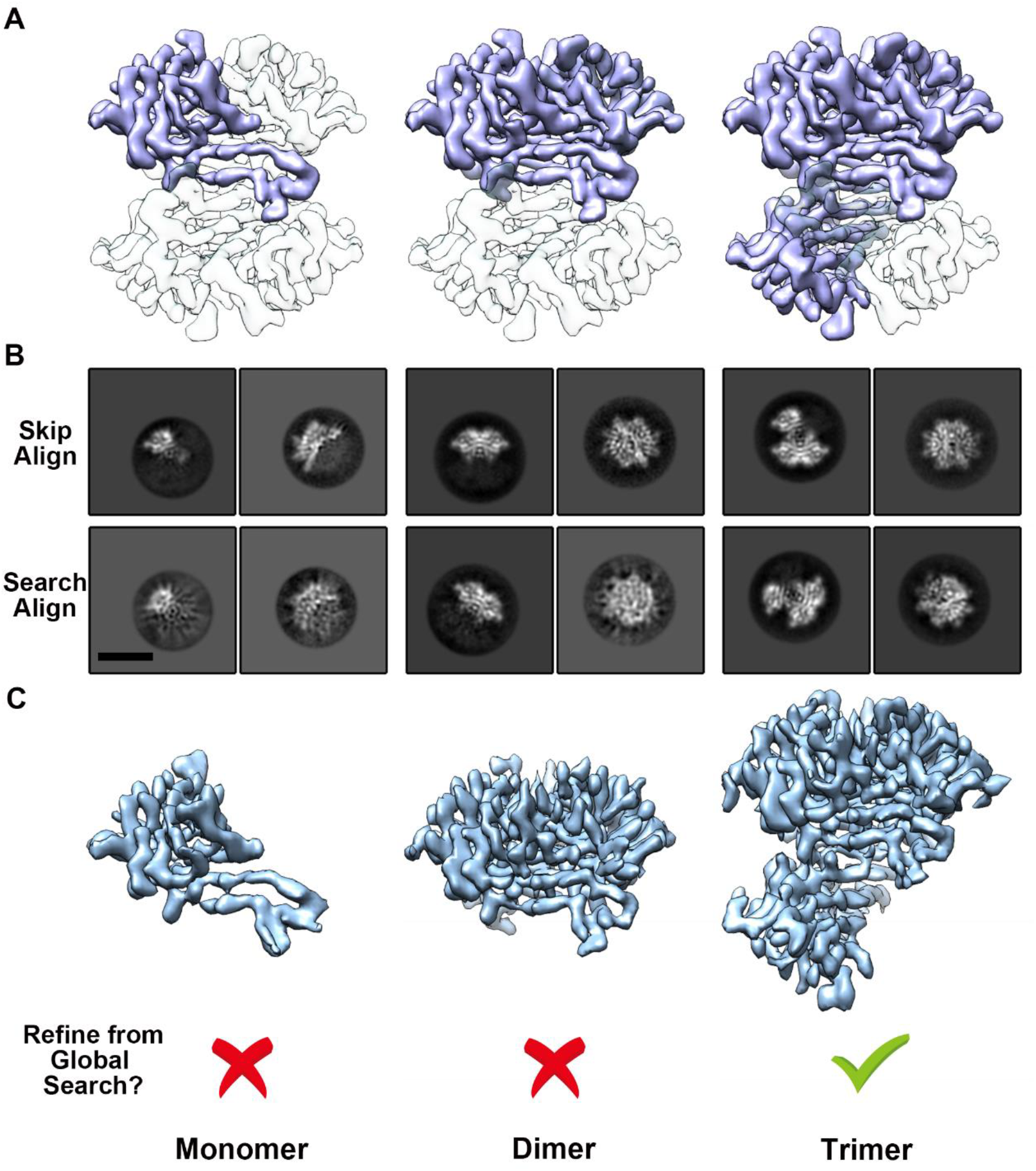
Reconstructions of subtracted SA in different oligomeric states. (A) The diagrammatic sketch of the subtraction in raw biotin-SA particles. The white parts were subtracted from individual particle images based on related angular information with the blue part left for image processing (monomer, dimer and trimer from left to right panel). (B) 2D classification results from subtracted datasets in different oligomeric states either using the angular information generated from the 3D refinement of SA tetramers (Skip Align in Relion) or omitting the angular information in a reference-free mode (Search Align in Relion). (C) The 3D reconstructions of the three subtracted datasets individually. The top row are the reconstructions using the angular information generated from the 3D refinement of SA tetramers. Bottom row indicates that only the trimeric dataset could perform a successful global refinement from scratch.

The 2D classification results of the three datasets were consistent with the 3D refinement, indicating that: 1) all datasets contained enough signal for reconstruction at high resolution if the angular information is correct; 2) the 39 kDa trimeric SA images already contained enough signal for image processing from scratch to obtain a high resolution structure. Meanwhile, it also indicated good 2D class averages with clear features would provide a high possibility of successful reconstruction. In our results, the 26 kDa dimeric dataset could already generate high quality 2D class averages of certain orientations. It might be the lack of accuracy alignment in other orientations that caused the failure in 3D refinement. We infer that the major constitutes of beta-strands in SA made the alignment difficult in some orientations. All the above results indicated a high possibility to obtain an asymmetric protein structure with molecular weight ~30 kDa at near-atomic resolution by SPA cryo-EM.

### Distribution of SA particles in the vitrified specimen

We noticed that even after the careful scrutiny of the SA particle images by 2D classification to remove all obvious junks or bad particles, there were only about 20% (79,289 vs 378,987 for the apo-SA, FigureS3) out of the seemingly good particles providing a correct high-resolution reconstruction after 3D classification. Indeed, no matter how hard we tried, the rest 80% particle images did not generate reconstruction with clear secondary structural details, even though they appeared very similarly in our eyes to the good particles in the high-resolution reconstruction. We also confirmed that the original micrographs containing these particles were of high quality. We were wondering what made the difference of the quality of the particles for their contributions to the high-resolution reconstruction. It has been hypothesized that the adsorption of protein molecules to the air-water interface could cause denature or partially unfolding of the protein^3,10,11^. We were wondering if the location of particles in the thin layer of vitreous ice caused the variation of the image quality for high resolution reconstruction. We therefore performed electron tomography of the same grid for single particle data collection of SA on graphene supporting film using VPP-Cs-corrector-coupled cryo-EM. 3D reconstructions of the tomograms were clear enough for us to depict the SA particle distribution in the specimen (Movie S2, S3, Figure 6 and S6). It is interesting to see that the SA particles distributed mainly in two different layers along the z-direction, one at the graphene-water interface (GWI) and the other at the air-water interface (AWI) (Figure 6A, Figure S6). There were very few particles between these two layers. This suggests that during the specimen preparation, the SA molecules either stuck to the graphene film or got adsorbed to the air-water interface. Surprisingly, particles at GWI had an uneven distribution, mostly in “clustering areas” (Figure 6A, red arrow) and only a few in “lacuna areas” (Figure 6A, blue arrow). This might be due to the effect of an uneven glow-discharge on the graphene surface. In contrast, particles on AWI showed a more uniform distribution (Figure 6B). Such a phenomenon was observed in both relatively thick (~50 nm, Figure S6C, Movie S2) and thin (~10 nm, Figure S6D, Movie S3) ice. The electron tomography analysis implied that the micrographs of SA single particles collected at zero-degree tilt actually reflected superposition of the particles at both GWI and AWI.

**Figure 6.**
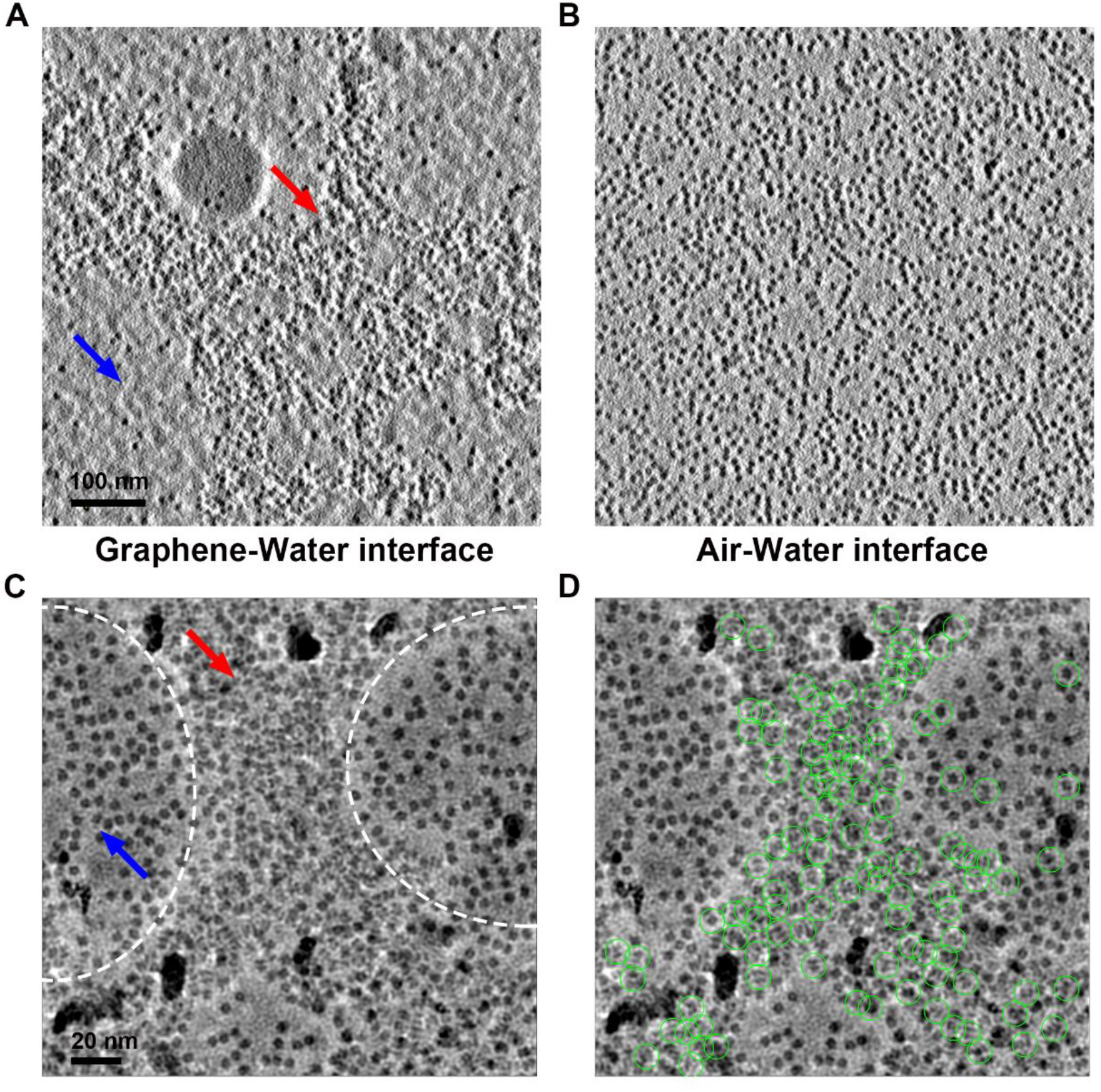
SA particle distributions on graphene-water and air-water interfaces. (A) The X-Y cross-section corresponding to the graphene-water interface from a reconstructed tomogram. The uneven distribution of particles are indicated as clustering area (red arrow) and lacuna area (blue arrow). (B) The X-Y cross-section corresponding to the air-water interface from the same reconstructed tomogram as in (A). (C) A micrograph containing clustering area (red arrow) from graphene-water interface and uniform distribution area from air-water interface (blue arrow). The boundaries of the two areas are marked by dashed lines. (D) The same micrograph as in (C), with the particles that contributed to the final high-resolution reconstruction circled in green.

We analyzed all the micrographs used for single particle reconstruction by sorting them by the percentage of good particles classified in the correct high-resolution reconstruction. We found that the best micrographs with the highest percentage of good particle images had an uneven distribution of the molecules (Figure 6C). The good particles that were classified in the correct high-resolution reconstruction came mostly from the clustering areas with a similar pattern to those on GWI as revealed by electron tomography (Figure 6D). The particles in the more uniformly distributing areas contributed very little to the final high-resolution reconstruction. Combining the cryo-electron tomography results and the single particle micrograph quality analysis, we may conclude that the SA particles attached to the graphene supporting film are better preserved for their native structures than those adsorbed to the AWI.

## Discussion

As SPA cryo-EM has become a powerful method in solving many supramolecular complexes with large molecular weight, people keep wondering how small a molecule can be solved at near atomic resolution by this method. Here, we demonstrated that using VPP and Cs-corrector, SPA cryo-EM can solve SA with molecular weight of about 50 kDa at ~3 Å resolution, good enough to determine the ligand binding site. By combining particle subtraction analysis, we could push the lower boundary of molecular weight further to at least a 39 kDa asymmetric tetramer at ~3 Å resolution. Although 3 Å resolution is not high enough to accurately assign every atom from a specific small molecule, with additional information of the possible conformations of the molecule, we could identify the binding pocket and possible interactions (like hydrogen bonds) between protein and ligands. More recent progress in algorithm development enabled the solution of macromolecules with large molecular weight and high symmetry at higher than 2 Å resolution^12,29,30^. Previous theoretical predictions suggested that SPA could determine the atomic resolution structure of protein with molecular weight as small as ~ 20-40 kDa ^31-33^. It is foreseeable that macromolecules as small as streptavidin or even smaller ones can be solved at high enough resolution to build atomic models of ligand *de novo*. Such a scenario would make the cryo-EM structure-based drug discovery more trivial. More importantly, the unique power of single particle cryo-EM in dealing with heterogeneous ligand occupancy and conformations in a single specimen could help accelerate drug screening process without any crystallization trials or crystal soaking, which is both time- and material-consuming. One may solve the structures of a target macromolecule co-existing with multiple ligand candidates in the same specimen and get multiple ligand-occupied states solved simultaneously within a few days of data collection and computation.

To our knowledge, this work is the first to solve a near atomic resolution structure of a protein smaller than 100 kDa with supporting film. Using single-crystalline graphene as supporting film may bring us the following benefits: 1) reducing the ice thickness (ice noise) without introducing a strong background noise (Figure S6C, S6D); 2) reducing the charging effect, radiation damage and local motions due to the graphene’s excellent electric and heat conductivity; 3) attracting and concentrating protein particles near GWI and reduce the adsorption to AWI. In practice, we found that the graphene supporting film also makes the cryo-EM specimen preparation more controllable and repeatable.

It was unexpected to found that SA particles in our frozen-hydrated specimens either stay on GWI or get adsorbed to AWI but very few stay in the bulk of the ice. This suggests that during the period of specimen preparation, the SA molecules separate quite fast into the two interfaces and do not come back into the liquid bulk once hitting the interfaces. Our results suggested that the particles stayed at the GWI were well-maintained for their high-resolution structural features while those adsorbed to the AWI were at least partially damaged. This may explain the difficulty of making good cryo-EM specimens with thin enough ice for many protein samples: the proteins quickly get to the AWI and become denatured or partially unfolded therefore making the specimen unusable. To avoid the proteins hitting AWI too fast, it may be helpful by reducing the Brownian motion rate. This could be achieved by reducing the temperature of the protein solution, increasing the viscosity of the solution, or increasing the thickness of the liquid layer over the holes on the EM grid. But these may all unavoidably reduce the contrast of the molecules in cryo-EM. An ultimate solution to prevent macromolecules from hitting the AWI is by blocking the AWI with either a supporting film such as the graphene or some inert surfactant that does not has any impact on the macromolecules’ structures as suggested by Glaeser^10,34,35^. Anchoring the macromolecules with certain affinity tags to the graphene or other electron-transparent supporting materials would be a possible solution. Only when we avoid the denaturation of macromolecules at the AWI, we may be take full advantage of single particle cryo-EM analysis in deciphering the distribution of molecular machines in its conformational landscape beyond the static atomic models.

## Materials and Methods

### Cryo-EM sample preparation

For biotin free apo-SA cryo-sample, 1 mg/ml commercially available streptavidin solution (New England Biolabs) was diluted to 0.2 mg/ml by 25 mM Tris-HCl buffer (pH 7.5, 75 mM NaCl). After centrifugation (12,000 g, 15 min), 4 μl diluted protein sample (0.2 mg/ml) was added to a pre-glow-discharged 300 mesh Quantifoil Au R0.6/1 graphene-coated grid in Vitrobot Mark IV (FEI Company) with 100% humidity and waited for 10s before blotting. The graphene-coated grid was prepared as previously described^24^. A blot force of -1 and blot time of 1s were applied to blot the grid after waiting. After blotting, the grid was plugged into pre-cooled liquid ethane.

For biotin-bound SA cryo-sample, the same dilution buffer as above containing extra 5 mM biotin (Sigma-Aldrich, St Louis, MO, USA) was used to prepare the protein sample (0.2 mg/ml streptavidin, 4 mM biotin). The biotin-SA solution was incubated on ice for 1 hour and then centrifuged at 12,000 g for 20 min. When preparing the cryo-sample, the waiting time before blotting was changed to 2s and blot force was changed to -2. The rest of steps were the same as above.

### Data collection on VPP-Cs-corrector-coupled EM

All the data were collected on the same 300 kV Cs-corrected Titan Krios microscope which is equipped with a FEI volta phase plate (FEI) by a K2 Summit direct electron detector with GIF Bio-Quantum Energy Filters (Gatan). After the cryo-specimens were loaded into the microscope, we first performed basic alignment of the microscope. Then we tuned the Cs-corrector and VPP at eucentric-focus with ~ -0.5 um defocus from eucentric-height at 195,000x magnification (TEM mode, micro-probe) as previously published procedure^22^. The microscope with a well-tuned Cs-corrector was then changed to EFTEM mode, and the low dose module exposure mode was set to nano-probe mode with 50 um C2 aperture at 215,000x magnification (EFTEM mode). The K2 detector was gain-corrected and Energy Filters was full tuned at the exposure condition. We have updated the previous version of AutoEMation so that it can perform fully automatic VPP data collection as well as VPP position change and initial phase-shift buildup for every ~40 images as previously established^22^. During the data collection period, the objective lens was set at eucentric-focus and the specimen was adjusted and imaged at a Z-position of -0.8 μm from the eucentric height for all the exposure holes within a 8-μm radius area. 32-frame super-resolution movies were collected in a 2.56s exposure time with a total dose of 50 e-/Å2 and super-resolution pixel size 0.26325 Å at the specimen level. In total, we collected 1,450 movie stacks for apo-SA in a 1-day session and 3,309 movie stacks for biotin-SA in a 2-day session.

We used SerialEM to collect VPP electron tomographic data on exactly the same apo-SA cryo-specimen for SPA data collection. Tilt series were collected from -54 degree to 54 degree with a 3-degree interval at 64,000x magnification (EFTEM mode, super-resolution pixel size 0.886 Å at the specimen level). For each tilt, the exposure time is 1.0s with 8 frames using a total dose 3.38 of 125 e-/Å^2^ in super-resolution mode.

### Image processing

Super-resolution raw frames of K2 camera were integrated to MRC format stacks by a local-written program Dat2MRC (developed by Bo Shen, unpublished). MotionCorr (-bin 2 -fod 4 -bft 200 -ssr 1 -ssc 1 -pbx 192) was firstly used for full-frame alignment and generated bin2-movie stacks for initial examination^4^. After initial examination of the movie stacks, the good uncorrected bin2-movie stacks were further processed by MotionCorr2 for a 5x5 patches drift correction with dose weighting (-PixSize 0.5265 -kV 300 -Iter 30 -Patch 5 5 -FmDose 1.56 -Bft 200 -Group 3)^36^. The summed bin4-images were generated with pixel size of 1.053 Å after the MotionCorr2 correction. The non-dose-weighted images were used for CTF estimation of defocus, astigmatism and phase shift parameters by Gctf^16^. The CTF fitting of each micrograph was examined and screened by checking the Thon ring fitting accuracy manually. The dose-weighted images were used for particle picking and reconstruction. For apo-SA dataset, 709,967 particles were automatically picked by Gautomatch (developed by Kai Zhang, http://www.mrc-lmb.cam.ac.uk/kzhang/Gautomatch/) from 1,385 micrographs. For biotin-SA dataset, 1,346,980 particles were picked by Gautomatch from 3,272 micrographs. After particles were extracted by Relion, a 120 Å high pass filter was applied to the particle stacks by relion_image_handler for a better 2D classification performance. The initial model was generated *de novo* by the 3D initial model in Relion using the SGD method. For each dataset, multiple rounds of 2D or 3D classification were performed in Relion to screen the best particles producing the two 3D reconstructions with D2 symmetry of apo-SA at about 3.3 Å resolution (23,991 particles) and 3.2 Å resolution (45,686 particles) based on the gold-standard FSC criterion. Additionally, a 3.1 Å resolution reconstruction could be generated by combing the two datasets together with final refined particles (69,677 particles in total).

For the asymmetric single particle analysis, the star files of related particles were extended four times by program relion_particle_symmetry_expand with D2 symmetry. Then the new star files were input into Relion for skip align 3D classification and particle subtraction following standard process.

For generating the subtracted datasets, the densities to be subtracted were manually adjusted in UCSF-Chimera^37^. Subtracted particles were re-refined with either local angular search (within 1.8 degree) or global search from scratch (initial 7.5 degree). For the global search refinement, initial models were generated from target apo-SA maps with 20 Å low pass filtering.

For tomography reconstruction, the tilt series raw stacks were firstly drift-corrected by MotionCor2. The fiducial free alignment and tomogram reconstruction was done by IMOD by standard procedure^38^. The final tomograms were generated with an eight-time binning (pixel size 7.088 Å) from super-resolution images.

### Model fitting and refinement

The atomic model of biotin-SA (PDB 1MEP) was fit into the EM density maps as a rigid body in UCSF-Chimera. The crystal structure fit well in the high-resolution EM density maps. Based on the map densities, we mutated and refined some side chains manually in Coot^39^ and run one round of real space refinement in PHENIX^40^.

## Acknowledgement

We thank Xiaomin Li and Tao Yang at the Tsinghua University Branch of the National Protein Science Facility (Beijing) for their technical support on the Cryo-EM and High-Performance Computation platforms. We thank Zhipu Luo at Soochow University for his help in atomic model refinement. This work was supported by grant (2016YFA0501100 to H.W.) from the Ministry of Science and Technology of China, grant (Z161100000116034 to H.W.) from the Beijing Municipal Science & Technology Commission.

## Supplementary Figures

**Figure S1.**
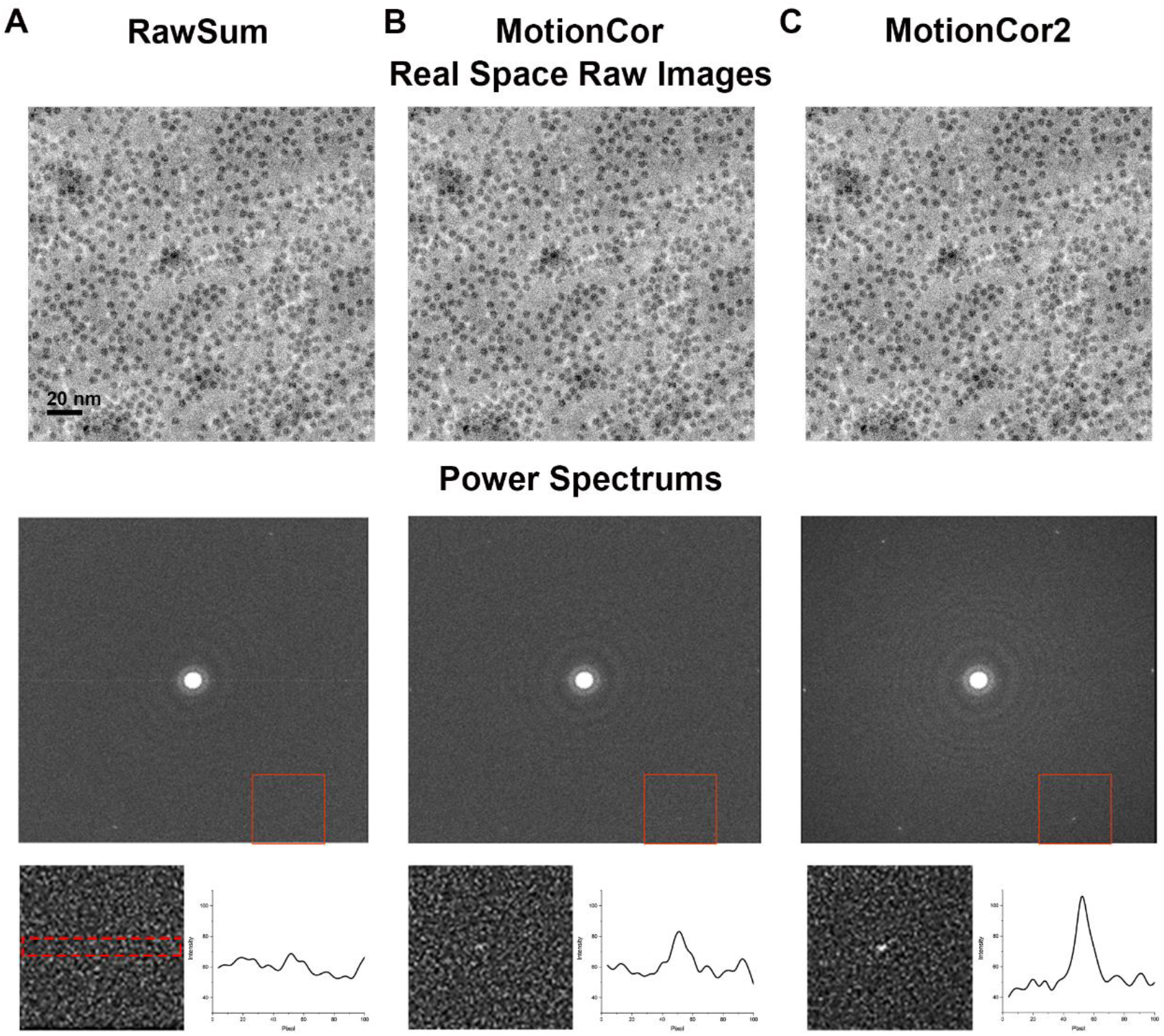
Motion correction of movie stacks of SA specimen on graphene supporting film. (A-C) The summation of movie stacks without motion correction, treated with MotionCor and with MotionCor2, respectively. In the top row are the summed images after correction. In the middle row are the corresponding power spectra calculated from the summed images. The reflections at 2.13 Angstrom resolution, reflecting the hexagonal lattice of graphene, are boxed in red. In the bottom row are the zoom-in of red squared region showing the reflection spot more clearly and the corresponding integrated intensity profile of each reflection within the red rectangular area as marked in the most left panel.

**Figure S2.**
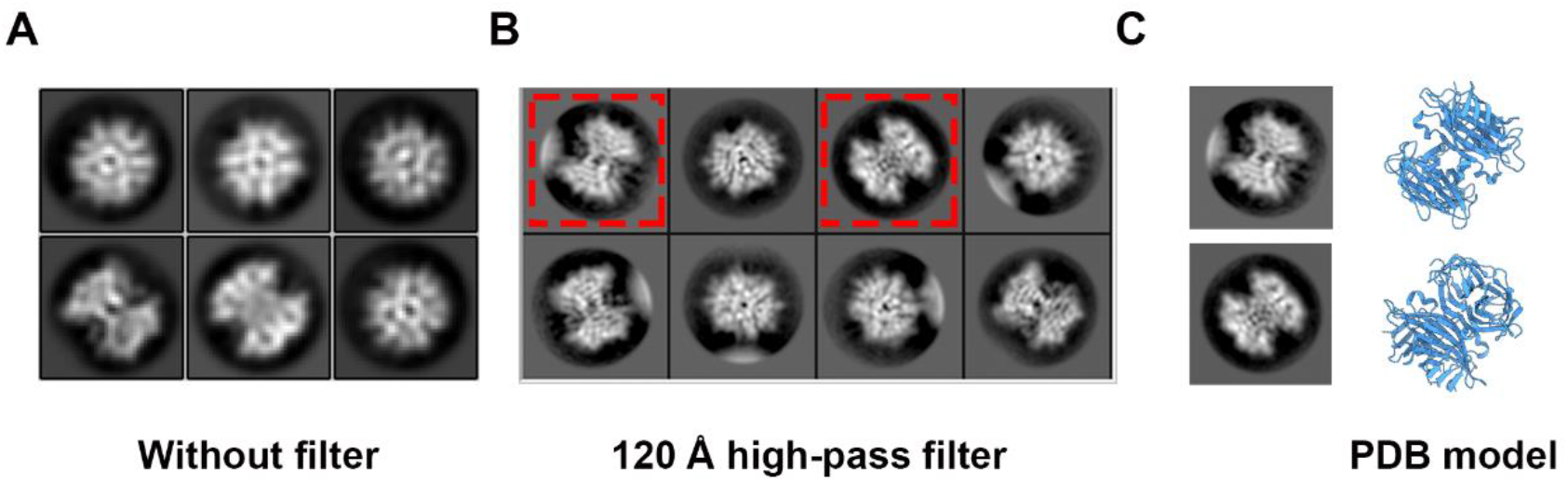
High pass filter for data processing. (A) Representative 2D class averages of SA particles without high-pass filter. (B) Representative 2D class averages of SA particles with high-pass filter at 120 Angstrom. (C) Selected 2D class averages as boxed in (B) in comparison with the similar views of atomic model of SA (PDB 1MEP).

**Figure S3.**
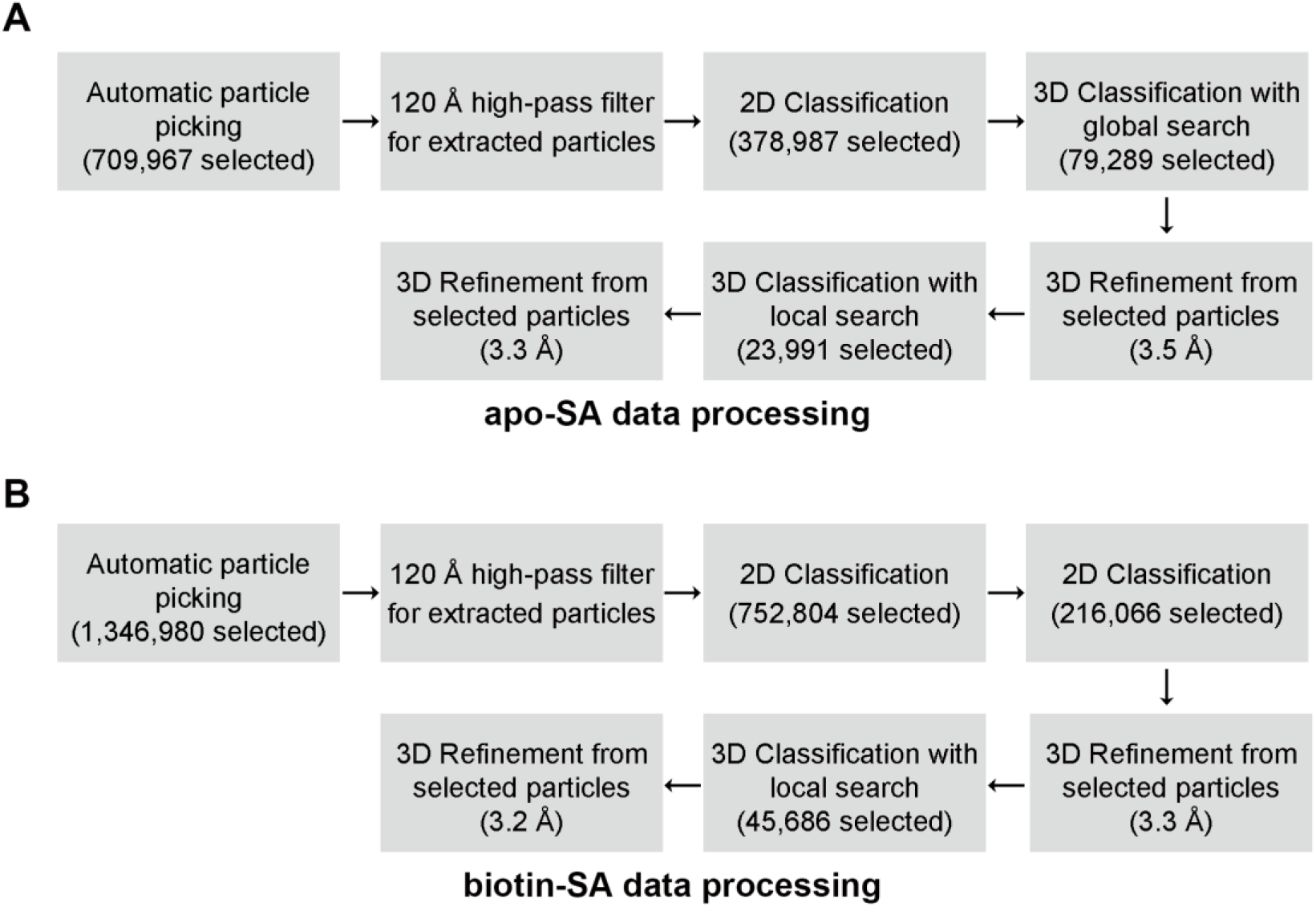
Single particle image processing flow charts. (A) Apo-SA image processing flow chart and (B) biotin-SA image processing flow chart.

**Figure S4.**
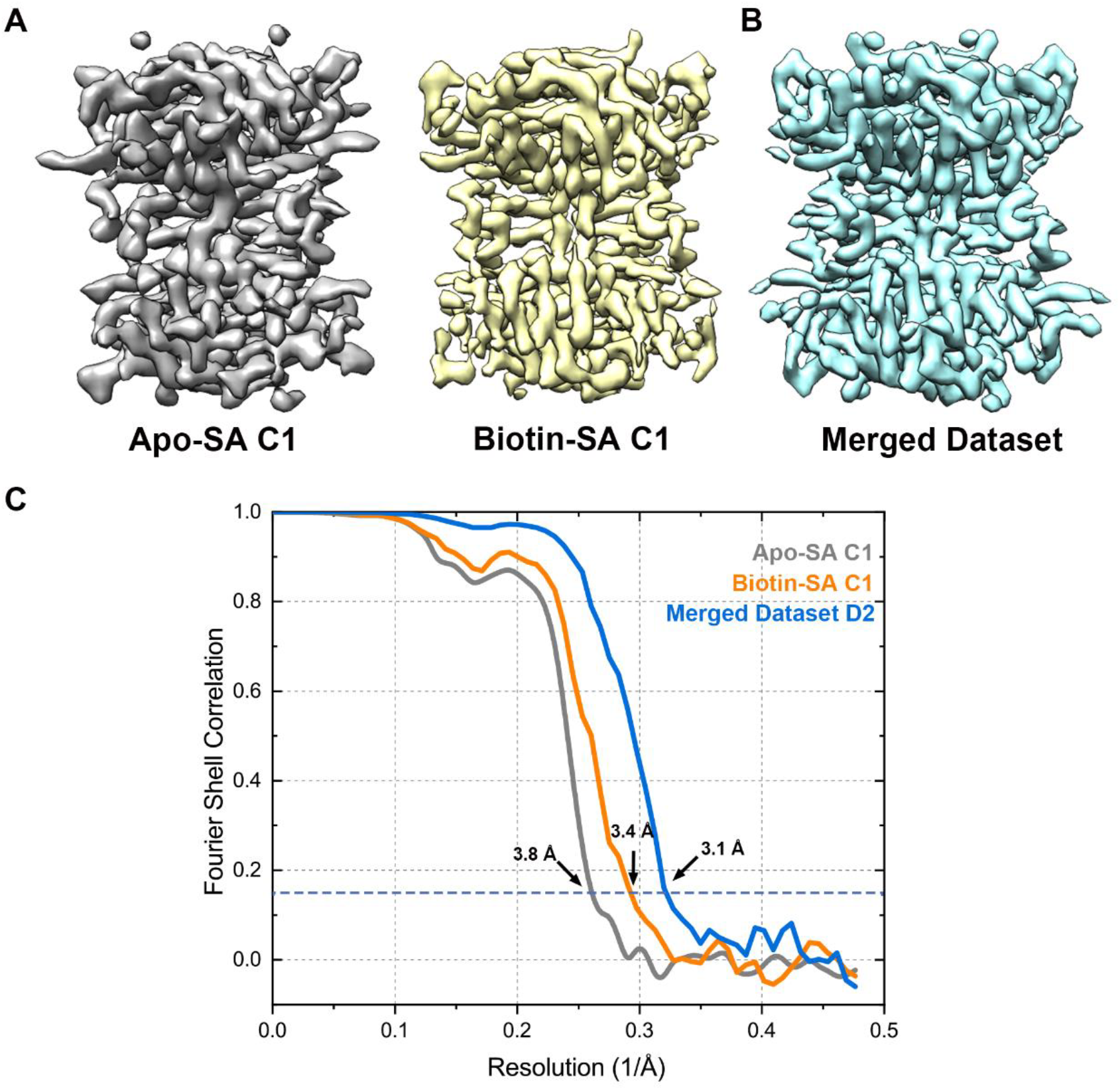
Reconstructions of the SA datasets with different symmetry imposition. (A) The C1 symmetry reconstructions of apo-state SA (3.8 Angstrom resolution) and biotin-bound SA (3.4 Angstrom resolution), respectively. (B) The reconstruction of merged-dataset (apo-SA + biotin-SA) with D2 symmetry imposition (3.1 Angstrom resolution). (C) The FSC curves of the three reconstructions with gold standard criteria.

**Figure S5.**
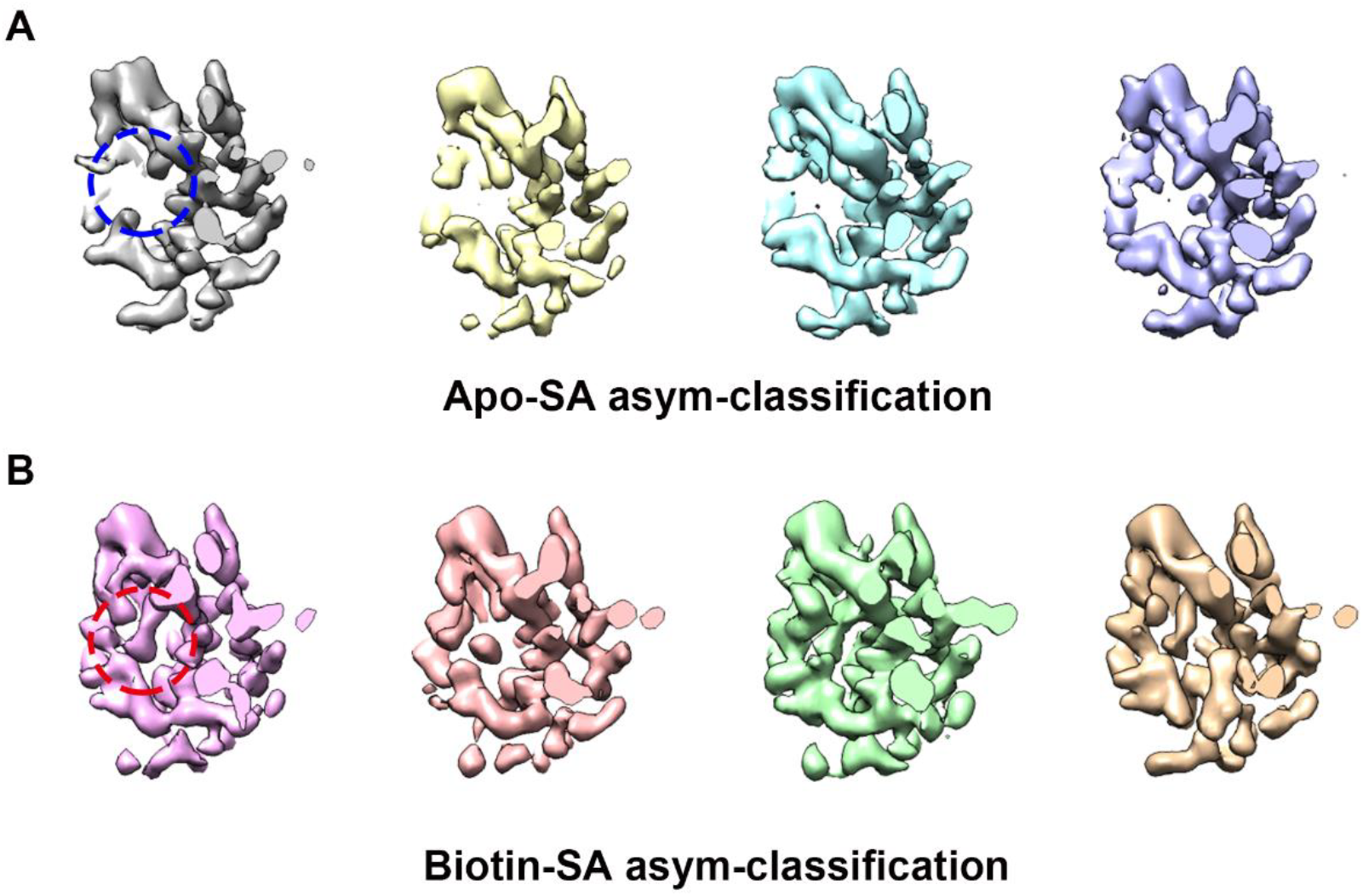
Asymmetric classification of homogenous dataset of SA. (A) Asymmetric 3D classification of the apo-SA dataset demonstrating empty biotin-binding pockets (blue circle) in all four classes. (B) Asymmetric 3D classification of the biotin-bound SA dataset demonstrating occupied biotin-binding pockets (red circle) in all four classes.

**Figure S6.**
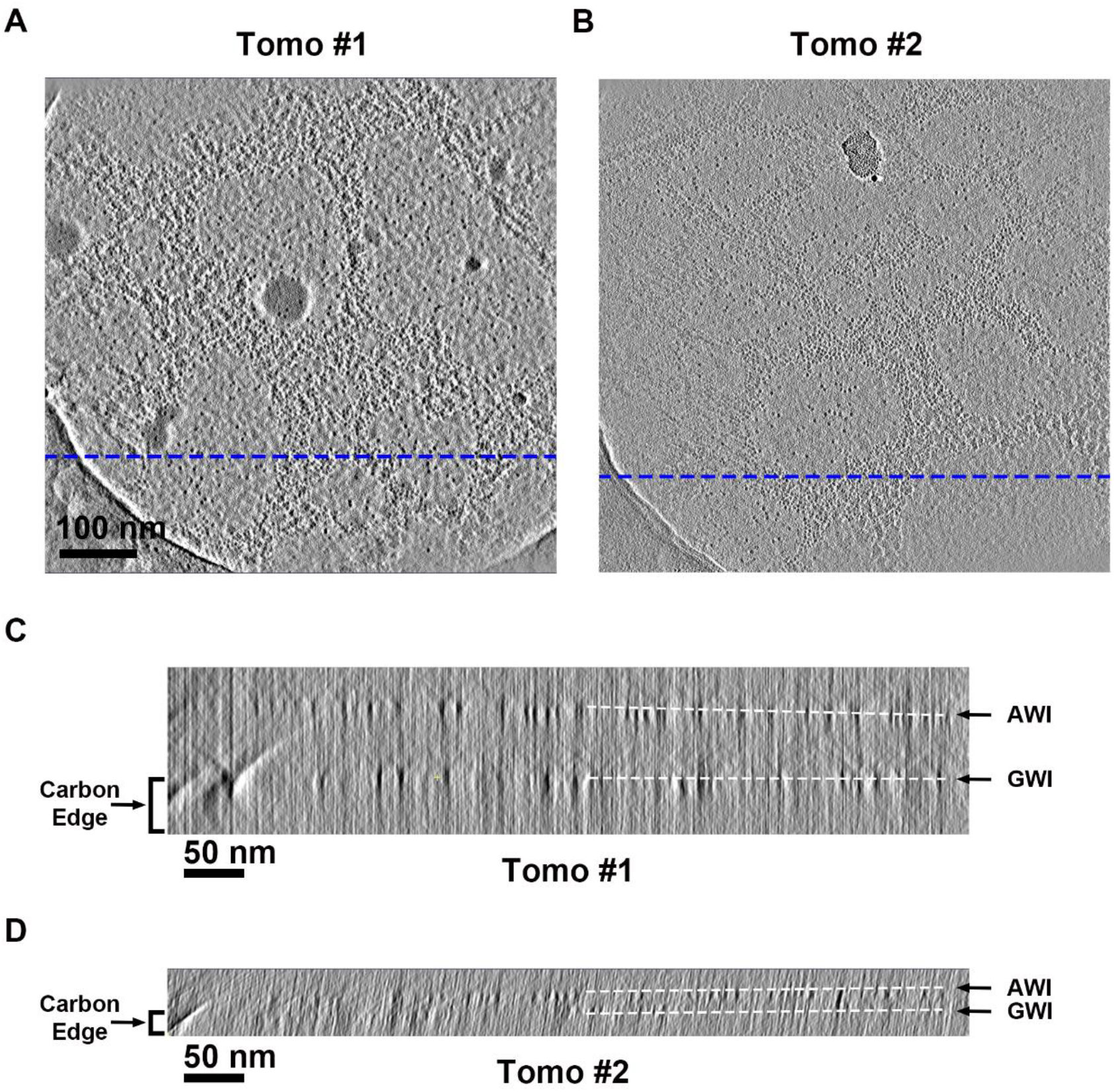
Cross-sections from electron tomographic reconstructions. (A) and (B) are the X-Y cross-sections of graphene-water interface from two electron tomographic reconstructions. (C) and (D) are the X-Z cross-sections from the blue dash lines plane in the tomographic reconstructions of (A) and (B), respectively. White dash lines in (C) and (D) indicate the air-water interface (AWI) or graphene-water interface (GWI). The location of carbon edge of the holes are indicated in (C) and (D).

## Supplementary Movies

**Movie S1. Fitting of atomic models in the cryo-EM densities of apo-SA and biotin-SA reconstructions**

This movie demonstrates the quality of the reconstructed apo-SA and biotin-SA maps in different areas with atomic models fitted. It also displays the major difference between biotin-SA (white) and apo-SA (green) reconstructions at the biotin binding pocket.

**Movie S2. Electron tomographic reconstruction of the apo-SA specimen in an area with thin ice**

This movie is related to Figure 6A-B, S6A and S6C, showing the reconstruction of a ~50 nm thickness vitreous apo-SA sample on graphene grid. The reconstruction shows the air-water interface (AWI), graphene-water interface (GWI) and the interlayer vitreous ice with different protein distribution behavior.

**Movie S3. Electron tomographic reconstruction of the apo-SA specimen in an area with thick ice**

This movie is related to Figure S6B and S6D, showing the reconstruction of a ~10 nm thickness vitreous apo-SA sample on graphene grid. The protein distribution in the area with ~10 nm thickness is similar to the ~50 nm area (Movie S2), but with a thinner interlayer vitreous ice.

